# Intranigral injection of Alpha-Synuclein pre-formed fibrils leads to BBB compromise and Bilateral Dopaminergic Neurodegeneration in A53T Alpha-Synuclein transgenic mice

**DOI:** 10.1101/2025.11.26.690780

**Authors:** Soumitra Ghosh, Han H. Lin, Sarah Chu, Hai Ngu, Nanzhi Zang, Renee Raman, Kimberly Stark, Oded Foreman, Somaye Hashemifar, Amy Easton, Baris Bingol, William J Meilandt

**Affiliations:** Department of Neuroscience, Genentech Inc. South San Francisco, USA; Department of Pathology, Genentech Inc. South San Francisco, USA; Department of Small Molecule Analytical Chemistry, Genentech Inc. South San Francisco, USA; Department of Artificial Intelligence, Genentech Inc. South San Francisco, USA

## Abstract

Parkinson’s disease (PD) is a progressive neurodegenerative disorder characterized by alpha-(α)-Synuclein neuronal aggregation and loss of dopaminergic (DA) neurons. Developing animal models that replicate PD’s neuropathological phenotypes is critical for understanding its pathophysiology and evaluating potential therapeutic targets. In this study, we show that direct unilateral injection of human α-Synuclein PFFs into the Substantia Nigra (SN) of mutant A53T α-synuclein overexpressing mice induce bilateral phosphorylated α-Synuclein (pS129) pathology in the SN. This pathology spreads to the striatum, cerebral cortex, and midbrain within 60 days and is accompanied by neuroinflammation in the midbrain and cerebral cortex. Additionally, we observed synuclein-dependent neurodegeneration, with a 50% reduction in Tyrosine Hydroxylase (TH) intensity in the SN and a 40% reduction in Striatum, both bilaterally. The model also revealed a compromised blood-brain barrier (BBB) and T-cell infiltration in the PFF injected animals, correlating with pS129 pathology and neuroinflammation. Taken together, we developed a mouse model that recapitulates multiple PD phenotypes, providing a valuable platform for testing therapeutic strategies targeting human α-Synuclein pathology and for exploring CNS-peripheral immune interactions in PD.

**Graphical Abstract:** 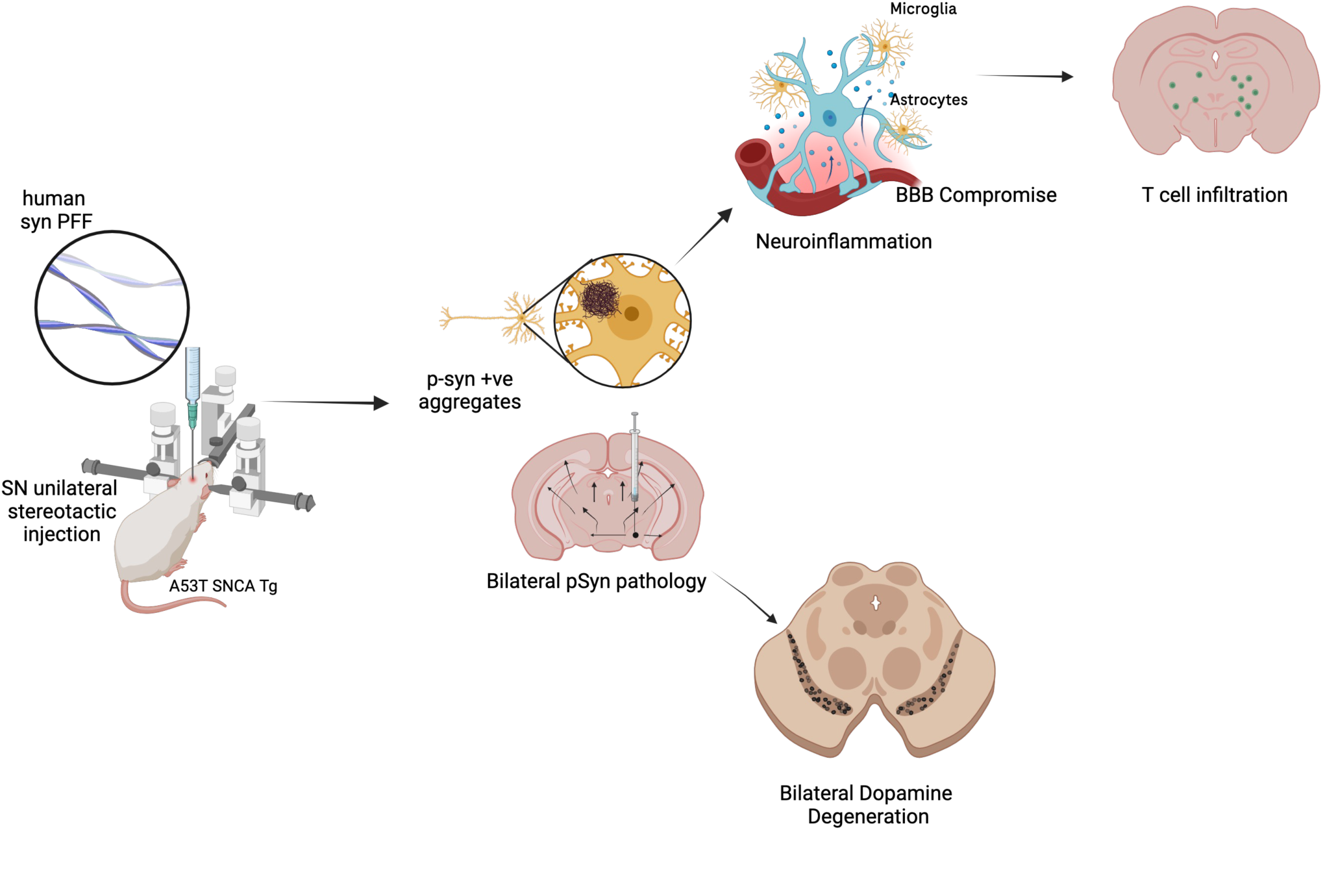

## INTRODUCTION

Parkinson’s disease (PD) is the second most common neurodegenerative disorder clinically diagnosed by the advent of motor and non-motor symptoms [1]. The signature hallmark of PD pathogenesis is the presence of Lewy bodies in neurons and loss of dopaminergic (DA) neurons in the Substantia Nigra (SN) [1–3]. Lewy bodies are proteinaceous cytoplasmic inclusions of aggregated α-Synuclein found in neurons throughout the brain. These inclusions are harmful to neuronal health and ultimately lead to neuronal death [1,3–5]. α-Synuclein pathology follows a characteristic neuronal migration as the disease progresses from different stages of severity, concomitant with behavioral deficits. This “spreading” feature suggests that α-Synuclein may form prion-like conformations that aid in the disease progression [2,4,5]. Genetic evidence strongly supports the role of the α-Synuclein gene (SNCA) in the etiology of PD, with gene duplications or triplications of wild-type SNCA[6], or rare mutations like A53T being identified as causative factors [7,8]. Currently there is no disease modifying therapy available for PD patients, but several clinical trials testing new therapeutic strategies are being pursued [9].

Transgenic (Tg) mouse models of synucleinopathies have been created to either overexpress human wild-type (WT) synuclein or familial forms of mutant synuclein (A53T or A30P), that develop synuclein inclusions, gliosis, and loss of neuronal TH staining predominantly in aged animals > 8 mo of age. [4,10,11]. For example, human α-synuclein overexpressing mice, known as Line 61, develop α-Synuclein inclusions at 12 months old and TH loss at 14-month age [12]. M83 mice overexpressing A53T synuclein mutation under the prion promoter develop cortical α-Synuclein inclusions as early as 8 months that spread to different brain regions with age. The animals develop motor neuron impairment and neuronal loss around 12-months-of-age, but specific dopaminergic loss in midbrain has not been reported [7]. A recent study overexpressing WT human α-Synuclein reported increased pS129 pathology and associated TH loss in the striatum, but not SN, at 13 mo of age along with BBB compromise [13]. Microvascular BBB alterations have also been reported in Thy1-aSyn line 61 transgenic mice at 6 months old [14]. These findings highlight the need for extensive aging for these models to develop dopaminergic neuronal degeneration.

To develop α-Synuclein inclusions in locations relevant to human PD patients, injection models have been highly adapted. Injection of pre-formed synuclein fibrils or virus overexpressing human synuclein or its mutants have shown tremendous potential. Synuclein overexpression in striatum or SN in WT mouse and rats have depicted the interaction of synuclein with dopaminergic neurons and its toxicity leading to loss of dopaminergic neurons [3,15–17]. The viral overexpression also leads to synuclein positive inclusions in regions of interest but do-not recapitulate the spreading nature of pathological synuclein from midbrain or striatum to other regions such as cortex as observed in later stages of PD in patients [18]. In the synuclein fibrils spreading model space, the first reports came when synuclein aggregates isolated from PD patient brains were able to spread to different regions of the brain when injected in striatum [19]. Thereafter, fibrils of synuclein have been developed in silico and used to study in-vivo. The majority of the studies have resorted to intrastriatal injection in the PFF model. Fibrils made from mouse synuclein when injected in WT mouse spread the synuclein pathology robustly to SN and other areas of the brain including cortex in the injected side. 35% loss of dopaminergic neurons has been reported in this model 180 days post PFF injection [19,20]. Similar spreading and synuclein associated pathology spreading have been reported in rats [21] but fibrils generated from human synuclein are less capable of spreading to other neurons in WT animals [22]. Injection of human synuclein fibrils in striatum of SNCA Tg mice on human synuclein transgenic background also show robust unilateral synuclein pathology, microgliosis and loss of dopamine transmitters [23]. Intrastriatal injections of human PFF in M83 mice led to bilateral spreading of synuclein inclusions in midbrain, striatum and cortex. Mortality was also observed in the PFF injected animals 75 days post injection [19]. In-silico modeling of mouse PFF injections in SN of WT mouse suggests a distinct pS129 spreading pattern when compared to intrastriatal PFF injections [24]. In published research from our group, we have observed a significant increase in neuroinflammation and pS129 pathology spreading in M83 mice 90 days post striatal injection. We have also shown that monomeric synuclein control striatal injections did-not develop any phenotype significantly different from saline injections [25,26].

In this study, we injected human synuclein PFFs directly in the Substantia Nigra (SN) of α-Synuclein A53T mutant overexpressing mice rather than using the traditional approach of injecting PFFs in the striatum. We observed a robust bilateral spreading of pS129 pathology in the midbrain, striatum and cortex. We also implemented an artificial intelligence (AI)-based workflow for TH intensity analysis, which provided a rapid and non-biased assessment of neuronal health in both the Substantia Nigra pars Compacta (SNpC) and Substantia Nigra Reticulata (SNR). A 50% loss of TH intensity was observed in SNpc and SNr depicting the effect of synuclein pathology on neuronal health. Additionally, we observed microglial inflammation, CD68 activation, and clustering of microglia throughout the brain attesting to the role of neuroinflammation in PD. To our surprise, there was significant BBB compromise and T cell infiltration in the brains of PFF-injected animals further recapitulating the additional PD associated phenotypes. This model effectively captures several PD-associated phenotypes at a single timepoint, allowing for in-vivo testing of therapies and investigation of rescue for multiple synuclein-associated phenotypes.

## METHODS

### Pre-formed human ɑ-Synuclein fibrils

Pre-formed human α-Synuclein fibrils were commercially sourced from Stressmarq Inc. (Cat #s SPR-317/ 322). The PFF aliquots of 5 mg/ml were stored in −80 °C. Prior to injection, the aliquots were thawed and sonicated for 5 mins at 30 sec on/off pulses in an enclosed bath sonicator (Qsonica, Q500). The PFF was examined under electron microscope (EM) post sonication. To perform EM, a 5 μL sample was applied to the continuous carbon EM grids (Electron Microscopy Sciences) and left for 120 seconds before blotting with Whatman #1 filter paper and dried fully. All TEM images were acquired using the Talos 200C microscope (Thermo Fisher Scientific Inc., Waltham, MA, USA) operating at 200 kV and equipped with a CetaD camera.

### Mouse husbandry and study design

All animal studies were reviewed and approved by the Genentech Institutional Animal Care and Use Committee (IACUC). All in-vivo studies were conducted in full compliance with regulatory statutes, IACUC policies, and NIH guidelines. The SNCA transgenic mice overexpressing the A53T mutation under prion promoter were obtained from JAX (A53T α-Synuclein transgenic line M83; JAX:004479) [7]. The animals were bred and maintained at Genentech on a C57BL/C3H background in a pathogen-free facility under standard conditions and provided with regular chow and water. Throughout the study, animals were maintained on a 12/12 light/dark cycle with a temperature between 22 ±2 °C and 50% relative humidity. A total of 24 animals were divided in three groups: 5 in the non-injected group (3 males, 2 females), 4 in saline (2 males, 2 females) and 15 in PFF (8 males,7 females) injected groups respectively. The animals were stereotactically injected at approximately 5.5 months old and sacrificed 8 weeks post PFF injection. Weights were monitored to check on the overall health and animals which showed immobility paired with hind limb paralysis or lost more than 15% body weight were sacrificed and tissues were collected for downstream analysis prior to the scheduled sacrifice date (Supple Fig 1 B).

**Figure 1.**
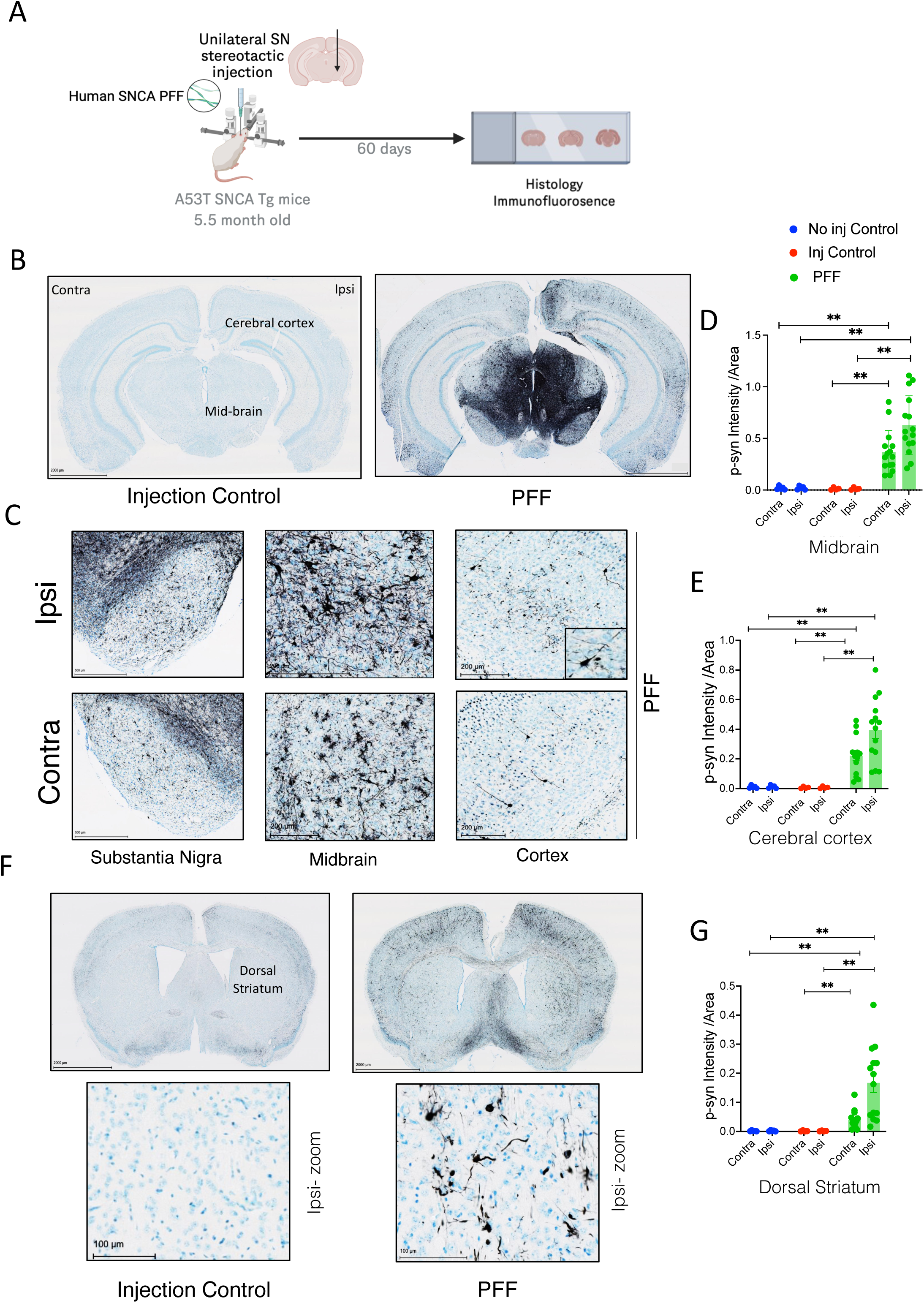
Unilateral SN PFF injection leads to bilateral spreading of pS129 associated pathology. **A)** Schematic representation of the in-vivo study; Created in BioRender. Ghosh, S. (2025) https://BioRender.com/b9z51j0 **B)** Representative brain sections covering midbrain and cerebral cortex depicting the pS129 Ab positive staining (black) in saline and PFF injected animals, Scale bar-2000 µm. Light Thionine Nissl (blue) for tissue area. **C)** Representative images for the PFF injected animals showing pS129 positive staining in Substantia Nigra, Scale bar-500 µm; Midbrain and Cortex, Scale bar-200 µm respectively. **D & E)** Quantification of p pS129 intensity normalized to segmented areas: midbrain and cortex (n= 5 non-injected, 4 saline, 14 PFF), Kruskal-Wallis test followed by Dunn’s multiple comparison test, **p-value <= 0.005. Asterisks (*) denote pairwise statistical significance. **F)** Representative brain sections covering dorsal striatum depicting the pS129 Ab positive staining (black) in saline and PFF injected animals, Scale bar-2000 µm. Light Thionine Nissl (blue) for tissue area. Lower panel shows magnified ipsilateral staining in saline and PFF injected animals, Scale bar-100 µm G) Quantification of pS129 intensity normalized to dorsal striatum area (n= 5 non-injected, 4 saline, 14 PFF), Kruskal-Wallis test followed by Dunn’s multiple comparison test, **p-value <= 0.007. Asterisks (*) denote pairwise statistical significance.

### Stereotactic Surgery

The animals were subjected to stereotactic surgery at approximately 5.5 months old. For stereotactic injections, the animals were anesthetized under isoflurane. A burr drill was used to create a small hole in the skull and a microinjection syringe was inserted to target the substantia nigra pars compacta unilaterally (anterior–posterior, +/- 3.0, medio-lateral, + 1.5, dorso-ventral, -4.6 from bregma). 2 μl of saline or PFF at a flow rate of 0.25 μL / minute [17]. Post-surgical incisional pain was treated with meloxicam.

### Behavioral activity

Mice were given an adaptation period of at least 60 minutes in their home cages within the testing room prior to the experiment. The rearing activity was measured using clear thermoplastic cages with dimensions of 40.5 cm in width, 40.5 cm in length, and 38 cm in height (San Diego Instruments). These cages were equipped with infrared (IR) beams set 3 cm above the floor to detect rearing behavior. The activity of the mice was recorded for a total duration of 30 minutes, starting from the time they were placed into the locomotor chambers. The testing took place under bright lighting conditions. Rotarod experiments were initially planned to assess motor coordination; however, PFF-injected animals developed severe and rapid hindlimb impairment near the mortality endpoint (Supple Fig 1B). This profound motor deficit compromised the animals’ ability to navigate the rotating rod, thereby precluding the collection of reliable data during late-stage pathology.

### Tissue processing

The mice were anesthetized and transcardially perfused with 30 ml of 4% PFA followed by 30 ml of 20 mM cold PBS (pH7.4) at a constant flow rate using a pump. The brains were then incubated with 4% PFA overnight (ON) followed by 20% glycerol and 2% Dimethyl Sulfoxide (DMSO) ON to prevent any freeze artifacts. Next, they were transferred to fresh 1X PBS containing 0.01 % sodium azide and sent to NeuroScience Associates Inc (NSA) for tissue sectioning. At NSA, the brains were embedded up to 25 per block and prepared for coronal sectioning within a gelatin matrix utilizing MultiBrain® Technology (NSA, Knoxville, TN)[25,26] . The brains were cured with a formaldehyde solution on a block and rapidly frozen by immersion in 2-methylbutane chilled with crushed dry ice and mounted on a freezing stage of an AO 860 microtome. The MultiBrain^®^ block was sectioned coronally with a setting on the microtome of 30 µm. Sheets of sections were stored in a cryoprotectant (30% glycerol, 30% ethylene glycol in PBS) at –20°C until subjected to immunostaining.

### Immunostaining and Imaging

For TH, pS129 and CD68 chromogen staining, the sheets of brain sections were stained as described on the NSA website and as described previously (https://www.neuroscienceassociates.com/ technologies/staining/; [25,26]). In short, pS129 a set of every twelfth 30 µm section (an interval of 360 microns) and TH and CD68 a set of every sixth 30 µm section (an interval of 180 microns) was stained in free-floating conditions. After a hydrogen peroxide treatment and rinses, each set of sections were immunostained with the primary antibodies as indicated in the table (Antibody List), ON at room temperature. Following rinses, a biotinylated secondary IgG antibody was applied. Vector Lab’s ABC solution Catalog # PK-6100 at a dilution of 1:222 was applied. All sections were treated with diaminobenzidine tetrahydrochloride (DAB) chromogen and hydrogen peroxide. The chromogen for the pS129 and CD68 stained sections included nickel (II) sulfate. Light Thionine Nissl counterstain was applied to αSyn-pS129 and TH slides prior to mounting. CD68 slides were counterstained with Neutral Red.

For CD3 DAB staining, serial sections of 30 µm covering the midbrain were stained in free-floating conditions. Sections were initially washed in PBS and then treated with 3% hydrogen peroxide in PBS. Following this, the sections were washed again with PBS and PBST and then blocked using 5% bovine serum albumin (BSA) in PBST. The primary antibody for CD3 was diluted in 1% BSA in PBST and the sections were incubated ON on a shaker at 4°C. On the following day, the sections were washed in PBST, incubated with biotin-conjugated secondary antibodies in an appropriate buffer, followed by treatment with an avidin–biotin–horseradish peroxidase (HRP) complex (Vector Laboratories #PK6100), and developed using Sigma diaminobenzidine tetrahydrochloride. The sections were then mounted and allowed to air dry ON. Finally, the slides were dehydrated in ethanol, cleared in xylene, and a coverslip was added using Permount.

All chromogen slides were scanned on a NanoZoomer S60 and a NanoZoomer XR whole slide imager (Hamamatsu, Bridgewater NJ) using a 20x (N.A. 0.75) objective lens at a scanning resolution of 0.46 μm/pixel.

For immunofluorescence-based tissue staining of Iba1, serial sections of 30 µm covering midbrain and cerebral cortex were stained in free-floating conditions. <u>BBB staining</u>: 4 animals in saline (2 males, 2 females) and 6 animals in PFF (3 males, 3 females) were randomly selected from the original PFF cohort for staining. Cy3IgG, Col IV, GFAP and CD31 were also stained similarly in 30 um brain sections covering ventral midbrain. IgG staining was performed on 30 um brain sections covering Striatum and SN along with the surrounding cerebral cortex. The sections were washed with PBS and PBST respectively, blocked for one hour in 5% Donkey Serum (DS) at room temperature. Post blocking, the sections were incubated ON at 4°C with the primary antibody (diluted as per the antibody table in <u>Methods</u>) in 0.5% DS. The following day, tissues were washed with PBST twice and sections were mounted with a gelatin solution and air-dried. A DAPI containing mounting medium was applied and overlaid with a cover slip. After drying ON, tissues were imaged under slide scanners or microscope. Immunofluorescence slides were scanned on an Olympus VS200 whole slide imager (Evident, Waltham MA) using a 20x (N.A. 0.8) objective lens at a scanning resolution of 0.274 μm/pixel for Cy3IgG and Zeiss Observer2.0 microscope was used to scan Col IV, GFAP and CD31 with a 10X or 20X (N.A. 0.8) objective lens. Maximum image projection was implemented on z-stack to capture the positively stained region within the 30 µm tissue section.

### Imaging data acquisition and analysis

An automated AI powered pipeline was developed at Genentech to segment Substantia Nigra pars Compacta (SNpC) and Substantia Nigra Reticulata (SNR) and measure the TH intensity within SNpC and SNR [27,28]. In short, the pipeline consisted of two neural networks (Supple Fig 4 A). The first neural network (YOLOv4) generated bounding boxes around the individual brain sections from a 5x5 tissue array of 25 TH-stained brain sections (processed and stained by NSA). The individual sections were cropped and resized to serve an input to the second neural network (U-Net) which segmented the SNpC and SNR region within the brain section. The centroid coordinate of the brain section was used to determine if the region belonged to the ipsilateral or contralateral side. A Matlab (MathWorks, Natick, MA) script using color thresholding and morphological operations was used to measure the TH positive area and intensity in the designated ROIs. The output was normalized to the ROI area and averaged across seven serial sections (Every twelfth 30-micron thick sections) spanning the SN for each animal. A modified pipeline has been adapted based on the same workflow to segment the striatum region (Dorsal Striatum) and measure the TH intensity in 4 serial 30-micron thick sections covering the striatum. To generate the manual segmentation data for the validation of AI based analysis, an expert in neuroanatomy of mouse brain manually drew ROI for SNR and SNpC in a subset of animals. The TH intensity was calculated in those ROIs using the Matlab script mentioned above. The efficiency of these models and their ability to measure the same area for an animal in both hemispheres was demonstrated by the Intersection Over Union (IOU) parameters (Supple Fig 4 B, C). We also selected a random subset of animals from our study, analyzed the TH intensity using traditional manual segmentation methods and compared it with the values obtained from the AI segmentation (Supple Fig 4 D, E). The results exhibited a strong correlation between both methods of measurement, further validating the robustness of the platform for automated segmentation and TH intensity analysis.

Phospho α-Synuclein 129 staining was quantified in midbrain, cerebral cortex and dorsal Striatum on a Genentech image viewer platform. Post manual ROI segmentation, the staining intensity was measured using a MATLAB script which determines the positive stain pixel area using adaptive thresholding. The output was then normalized to the ROI area and averaged across six to seven 30 µm sections for each animal. Similar analyses were performed for CD68 staining in the midbrain and cerebral cortex region.

CD3 positive T cells were measured by first drawing a ROI to highlight the left and right midbrain. The cells were counted manually on ImageJ using a cell counter and normalized to the area of the ROI. Injection tract adjacent areas and meninges were excluded from the counting of T cells. An average of three 30 µm sections per animal was quantified as the output.

Fluorescently labeled Iba1-stained 30 µm sections were analyzed using Matlab. A YOLOv4 neural network was used to crop individual brain sections. Whole brain tissue area covering the cerebral cortex and midbrain was segmented using thresholding and morphological operations. Five sections in each animal were analyzed. Iba-1 positive staining was detected using a top-hat filter and post-processed using morphological operations. A size threshold of greater than 53 sq microns was used to measure Iba-1 cluster area. Iba-1 intensity and cluster area were normalized to the whole brain tissue section area.

The Collagen IV (Col IV) and GFAP positive staining was quantified by segmenting an arbitrary equal size and location area across all animals covering the ventral midbrain at 10 X zoom (GFAP) and 20X zoom (Col IV). Average values from three 30 µm sections per animal were analyzed. Captured images were converted to 8-bit and Integrated intensity (CFCU) was measured on ImageJ for the specific ROI and the output is an average of three sections per animal. GFAP and Col IV integrated intensity within the ventral midbrain ROI was measured and normalized to the area of the ROI. The IgG positive staining was quantified by measuring the IgG intensity in the ventral midbrain. Integrated intensity was measured on ImageJ for the specific ROI and the output is an average of three 30 µm sections per animal. The images were then converted to 16 bit and thresholded by MaxEntropy method on ImageJ.

All statistical analyses were performed on GraphPad Prism using either Student’s *t* test, followed by Welch’s correction and Kruskal-Wallis non-parametric test followed by Dunn’s multiple variable test. Correlation analysis was performed by simple regression analysis on GraphPad Prism software. Each data point on the graphs represents an individual animal. The significance level and the number of animals analyzed for each test are detailed in the Figure Legends.

**Table 1:** Antibody list.

| Antibody | Catalog # | Source | Dilution | Secondary Antibody Cat# | Secondary Antibody (Source) |
| --- | --- | --- | --- | --- | --- |
| Tyrosine Hydroxylase (DAB) | P40101 | Pelfreez | 1:24K | BA-1000 | Vector |
| phospho Synuclein 129 (DAB) | ab51253 | Abcam | 1:150K | BA-1000 | Vector |
| CD68 (DAB) | 600-401-R10 | Rockland | 1:50K | BA-1000 | Vector |
| IBA1 (IF) | 011-27991 | Wako FujiFilm | 1:1K | A21432 | Thermo Fisher |
| Cy3 IgG (IF) | 115-165-003 | Jackson ImmunoResearch | 1:400 | NA | NA |
| CD3 (DAB) | MAB4841 | Novus Biologicals | 1:7.5K | NA | NA |
| GFAP (IF) | 13-0300 | Invitrogen | 1:1K | A78947 | Thermo Fisher |
| Col IV (IF) | 2150-1470 | Novus Biologicals | 1:1K | A-21206 | Thermo Fisher |
| CD31 (IF) | AF3628 | Bio-Rad Lab | 1:1K | A-11057 | Thermo Fisher |

NA- Not applicable

## RESULTS

### SN targeted PFF injection induces p-S129-associated pathology and its subsequent spreading across the brain

Injection of α-synuclein PFFs directly into the brain of mice is a commonly used model to study the seeding and spread of α-Synuclein pathology in-vivo [25,26]. Specifically in this study, we injected 10 ug of human α-Synuclein PFFs (Supp Fig 1A) or saline, unilaterally into the SN of A53T SNCA transgenic mice and monitored them for 60 days (8 wks). During the study, PFF-injected mice lost body weight, a reduction in the rearing behavior at 5 and 8 wks post-PFF injection and an increase in hind-limb paralysis suggesting that α-Syn fibrils can impact neuronal function resulting in behavioral deficits when injected into the SN. (Supp Fig 1B-D). We and others have previously shown [19,25,26] that intrastriatal injections of hPFFs in A53T mice showed similar behavioral symptoms and reduced lifespan but those observations were made at 80 days post injection whereas in our current model the phenotype is accelerated and observed at 60 days post hPFF injection. These observations suggest that the pathogenic spread of hPFFs are much more rapid when injected into the Nigra than Striatum.

Examination of coronal brain sections found that PFF injections into the SN resulted in a significant increase in pS129-associated α-Synuclein pathology bilaterally throughout the brain (Fig 1 A-E). Compared to non-injected or saline injected controls, cell bodies and neuronal processes in the Substantia Nigra, Midbrain and Cortex of PFF injected mice showed marked increase in pS129 staining (Fig 1C, Supple Fig 2A). Quantitative analysis of the p-S129 intensity per area in the Midbrain (Fig 1D) and Cerebral Cortex (Fig 1E) showed significant elevations compared with control mice. Significant spreading of pS129 was also observed in the dorsal striatum of PFF injected animals compared with control mice (Fig 1F & G). pS129 staining was elevated in both the ipsilateral and contralateral sides of the brain, but more pronounced elevation was observed in the ipsilateral injected side. Additionally, we observed pS129 spreading in distal brain regions in the PFF injected animals (Supple Fig 2B). This observation aligns with prior findings and further confirms the ability of hPFF to spread throughout the brain post unilateral SN injection [15,19].

**Figure 2.**
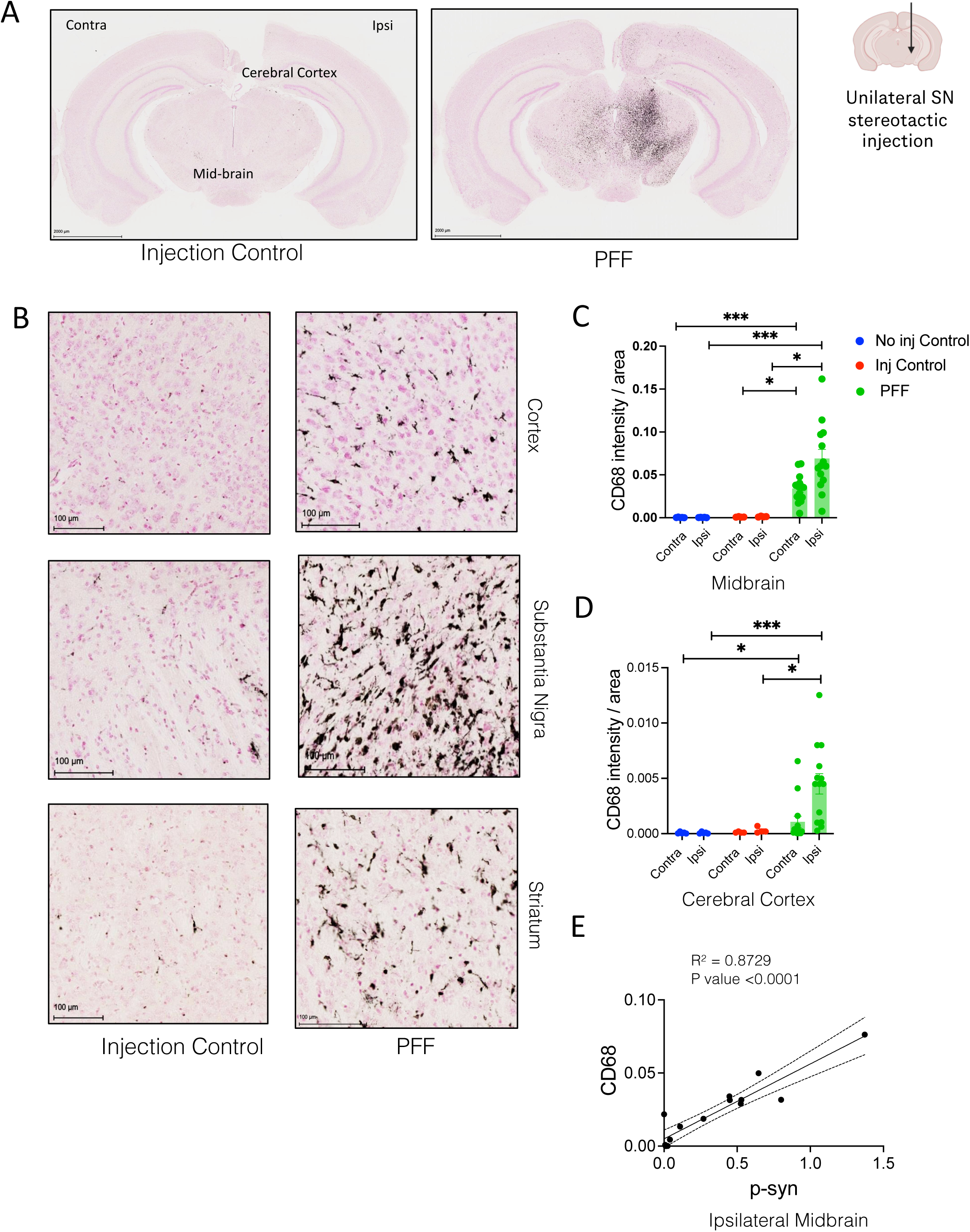
Unilateral SN PFF injection leads to neuroinflammation in the brain. **A)** Representative images covering midbrain and cerebral cortex depicting the CD68 positive staining (black) in saline and PFF injected animals. Neutral red (pink) staining for tissue area, Scale bar-2000 µm. **B)** Representative images for the positive staining for CD68 in Cortex, Striatum and Substantia Nigra in saline and PFF injected animals, Scale bar-100 µm. (I). **D, E)** Quantification of Cd68 intensity normalized to segmented Midbrain and Cerebral cortex area (n= 5 non-injected, 4 saline, 14 PFF), Kruskal-Wallis test followed by Dunn’s multiple comparison test, *p-value <= 0.05, ***p-value <= 0.0008. Asterisks (*) denote pairwise statistical significance. **F)** Simple regression analysis to plot the correlation between pS129 staining and CD68 in the ipsilateral midbrain of PFF injected animals.

### Pronounced microgliosis is observed in the brain post PFF injection

Neuroinflammation, specifically microgliosis, is now recognized as a key phenotype in PD pathogenesis [29]. Activated microglia has been observed in PD autopsied brains and in animal studies following α-Synuclein PFF injection, α-Synuclein AAV overexpression, and toxin (MPTP, 6-OHDA) injections that generate PD like phenotypes [30–34]. To determine if the synuclein pathology observed in the brains of hPFF injected mice were associated with altered neuroinflammation, we stained sections for Cd68, a lysosomal protein and marker for activated microglia [26] (Fig 2A). There was a significant elevation in CD68 positive microglial cells within the Substantia Nigra, Cortex and Striatum (Fig 2B) in the PFF-injected mice compared with controls. CD68 immunostaining was most prominent in the injected side of the brain, but was also elevated in the contralateral regions, consistent with the spreading of pS129 pathology to the same regions (Fig 2 D & E). In addition, there was a strong correlation between CD68+ neuroinflammation and α-Synuclein-associated pathology in the midbrain of PFF injected mice (Fig. 3F). To further confirm PFF-induced microgliosis, we measured IBA1 intensity in the cerebral cortex and midbrain and found a significant increase in PFF injected mice compared with non-injected and saline controls (Fig S3 A-F). Microglial cells display a ramified morphology and form clusters when activated. We found a significant elevation in microglial clusters in PFF injected mice compared with non-injected and saline controls (Fig S3 D-F). Together, these results suggest that nigral injections of α-Synuclein PFFs induced elevation in pS120 pathology and associated neuroinflammation.

**Figure 3.**
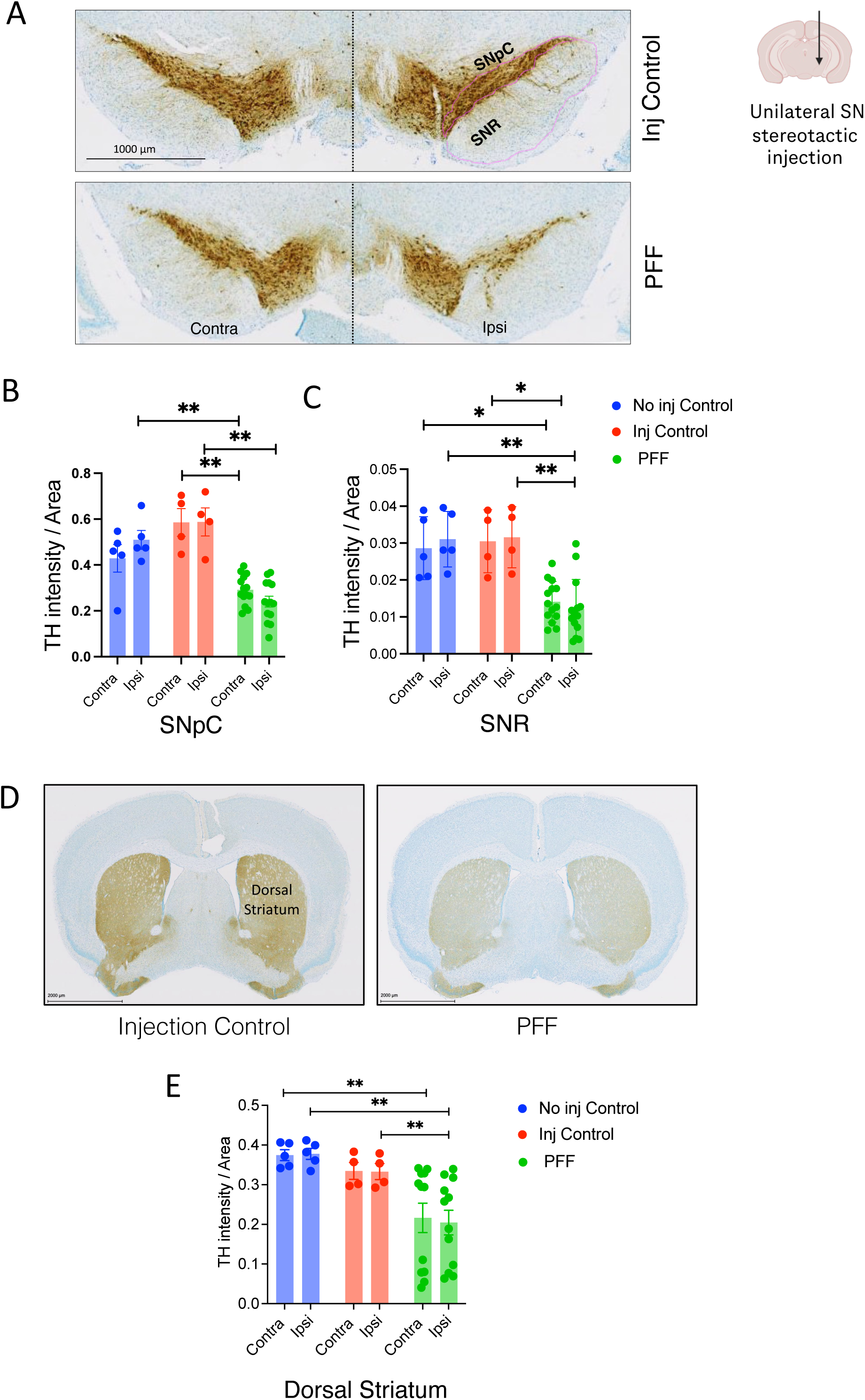
Unilateral SN PFF injection leads to bilateral loss of Tyrosine Hydroxylase intensity in the Substantia Nigra and Striatum. A & D) DAB staining for TH (brown) in the SN (A) and Dorsal striatum (D) of saline and PFF injected animals. Light Thionine Nissl (blue) for the tissue area of Saline and PFF injected animals. SN Scale bar-1000 µm. Striatum Scale bar-2000 µm. B, C & E) Quantification of SNR, SNpC and dorsal Striatum TH intensity as indicated in y axis normalized to ROI area (n= 5 non-injected, 4 saline, 14 PFF), Kruskal-Wallis test followed by Dunn’s multiple comparison test, *p-value <= 0.01, **p-value <= 0.009. Asterisks (*) denote pairwise statistical significance.

### PFF injection in the SN leads to loss of Tyrosine Hydroxylase in the Substantia Nigra

Dopaminergic neurodegeneration is a classical hallmark of PD. Elevations in pS129 and aggregated α-Synuclein are thought to contribute to neuronal dysfunction and loss [3]. To determine if the elevated pS129-α-Synuclein pathology caused dopaminergic neurodegeneration, we stained for Tyrosine Hydroxylase (TH) in brain sections of treated mice. Loss of TH immunostaining is often used as a measure for dopaminergic neurodegeneration in preclinical mouse models and in clinical samples [1,35,36]. We found a significant depletion of TH intensity following PFF injection compared with controls, in the SN and Striatum (Fig 3).

To facilitate the histopathological examination of TH loss in the SN, we employed an automated AI platform developed in-house [27]. Briefly, the AI model (Supple Fig 4A-D) first segments the region of interest for Substantia Nigra— reticulata (SNR) and pars compacta (SNpC), then quantifies the TH intensity within SNpC and SNR. Using this method, we found that there was a significant reduction in TH+ intensity (∼50%) in PFF-injected mice within the SNpC (Fig 3B) and SNR (Fig 3C), compared with control mice indicating dopaminergic neurodegeneration. We also observed a significant reduction (∼40%) in TH intensity on the ipsilateral and contralateral sides of the striatum (Fig 3D, E), likely representing loss of TH+ axonal projections from the SN. These results suggest that direct unilateral injection of synuclein hPFFs into the SN can induce bilateral dopaminergic neurodegeneration within 60 days post-injection in A53T overexpressing mice.

### PFF injection leads to T cell infiltration and BBB compromise in the brain

There is increasing evidence that alterations in the innate immune system may play a role in PD [37,38]. It is believed that neuroinflammatory cells, such as microglia, interact with peripheral immune cells to exacerbate PD pathology [39] . Studies in post-mortem PD brains have shown infiltration of T cells in brain parenchyma [30,40–42]. This phenomenon has also been observed in animal models of PD, particularly in those involving AAV mediated overexpression of WT or mutant forms of α-Synuclein [43,44]. In WT mice, T cell infiltration has been reported 5 months post mouse PFF injections [45]. Tg mice, specifically Line 61 SNCA mice, exhibit T cell infiltration at 11 months old without PFF injection and at 6 weeks following human PFF administration in adult mice [46,47]. Here, we investigated whether hPFF injections in SN could lead to T cell infiltration in A53T SNCA mice. To test this, we immunostained for the T-cell specific marker CD3, and observed a significant increase in CD3+ cells in the hPFF-injected mice compared with saline and non-injected controls. CD3+ cells were detected in the midbrain, hippocampus and cortex (Fig 4A-C, Supple Fig. 5A, B). The low counts of CD3 positive T cells on the ipsilateral injected side of the saline injected mice suggest that the surgical procedure of intra-nigral injections do not, on their own, cause T-cell infiltration. We observed a positive correlation between T cell infiltration and pS129 positive pathology along with CD68+ve neuroinflammation in the injected side of the midbrain (Fig. 4D, E). These observations suggest a potential crosstalk between microglial activation, pS129 pathology and T cell infiltration in this model.

**Figure 4.**
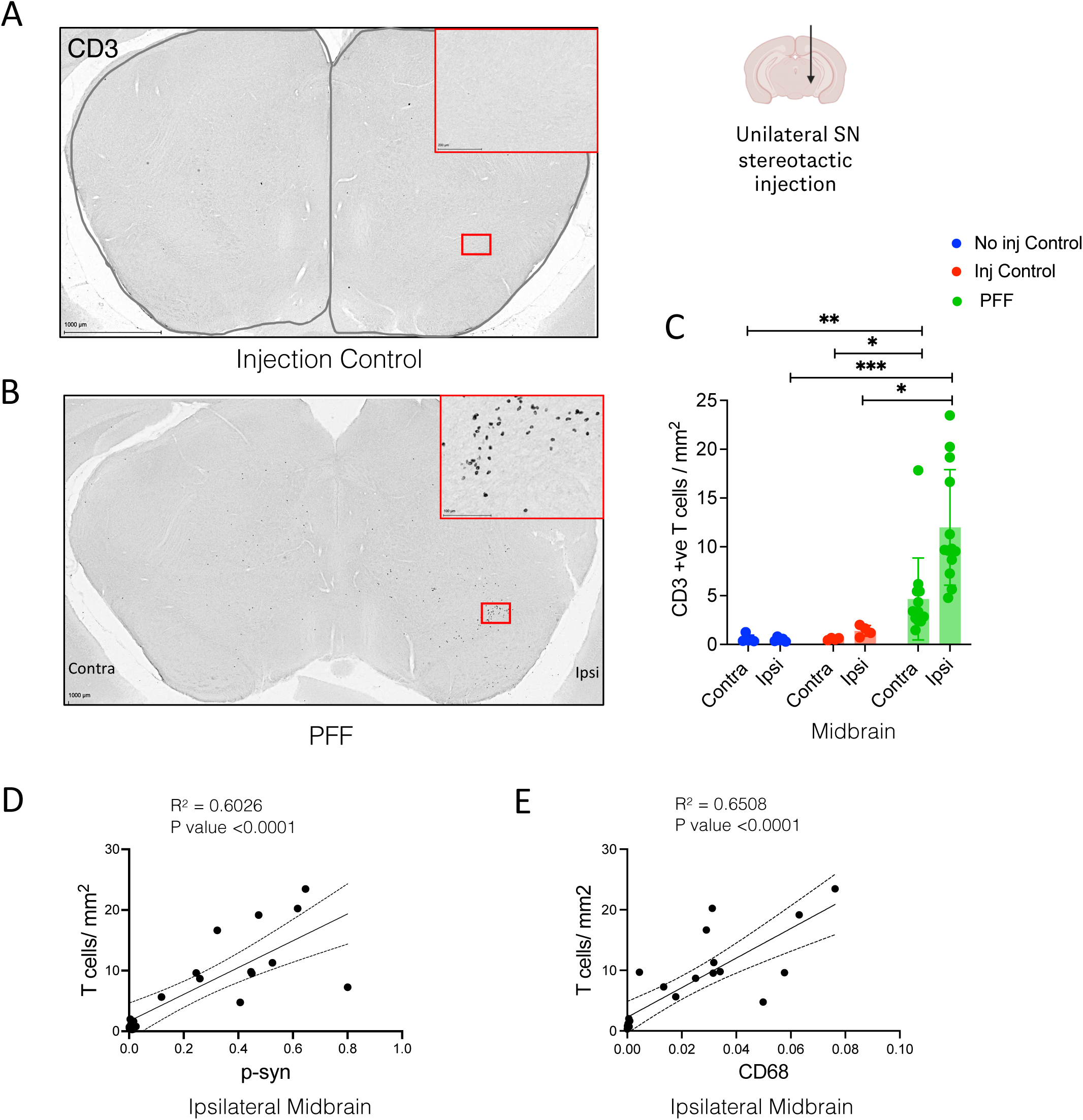
Unilateral SN PFF injection leads to T cell filtration in the midbrain. **A & B)** Representative images depicting the CD3 positive T cell immunostaining in saline and PFF injected animals. The areas segmented in gray line in the saline animal were used for quantification of CD3 positive signal, Scale bar-1000 µm. Red boxes represent the magnified images for the CD3 positive staining in the Substantia Nigra. **C)** Quantification of CD3 positive T cells per mm^2^ in the midbrain (n= 5 non-injected, 4 saline, 14 PFF), Kruskal-Wallis test followed by Dunn’s multiple comparison test, *p-value <= 0.02, **p-value <= 0.001, ***p-value <= 0.0005. Asterisks (*) denote pairwise statistical significance. **D & E)** Simple regression analysis to plot the correlation between T cells/mm^2^ and pS129 positive staining (D) and between T cells/mm^2^ and CD68 positive staining (E) in the midbrain of PFF injected animals.

**Figure 5.**
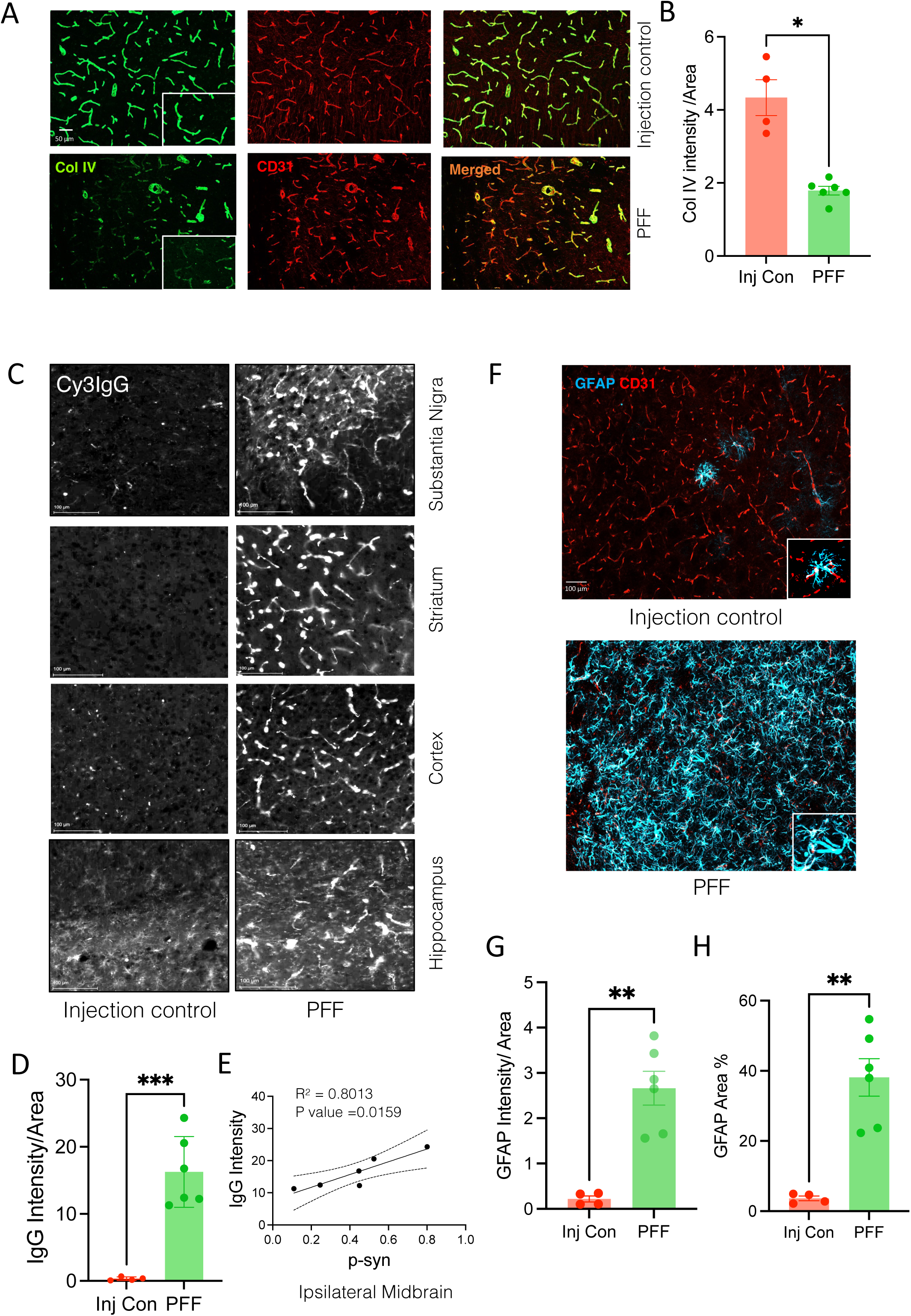
Unilateral SN PFF injection leads to BBB compromise and astrocyte migration in ventral midbrain. **A)** Representative images depicting the Col IV (green) and CD31 (red) immunofluorescence in the midbrain of saline and PFF injected animals. Scale Bar-50 µm **B)** Quantification of Col IV normalized to the ROI area (n= 4 saline, 6 PFF), Student’s t test with Welch’s correction comparison test, *p-value =0.0115. Asterisks (*) denote pairwise statistical significance. **C)** Representative images depicting the IgG positive immunofluorescence (white) of Cy3 conjugated IgG Ab in SN, Striatum, cortex and Hippocampus of saline and PFF injected animals, Scale bar-2000 µm **D)** Quantification of IgG intensity normalized to the ventral midbrain area (n= 4 saline, 6 PFF), Student’s t test with Welch’s correction comparison test, ***p-value =0.0004. Asterisks (*) denote pairwise statistical significance. **E)** Simple regression analysis to plot the correlation between IgG intensity and pS129 positive staining in the ipsilateral midbrain of PFF injected animals. **F)** Representative images depicting the GFAP (cyan) and CD31(red) positive immunofluorescence in ventral midbrain of saline and PFF injected animals, Scale bar-100µm **G)** Quantification of GFAP intensity normalized to the ROI area and GFAP positive Area % within fixed ROI (n= 4 saline, 6 PFF), Student’s t test with Welch’s correction comparison test, **p-value =0.001. Asterisks (*) denote pairwise statistical significance.

Our findings are consistent with those showing increased microglial activation and infiltration of peripheral immune cells following striatal injections of human or mouse PFFs into rodents [45,48]. It is unclear exactly how this increase in peripheral immune cell infiltration occurs. However, considering BBB leakage has been observed in PD patient tissues [49–51], and in aged BAC transgenic mice expressing WT a-synuclein [13], one possibility is that PFF injections could weaken the BBB thereby allowing infiltration of T cells into the brain parenchyma. To test this, we performed immunostaining for Collagen IV, the most abundant protein in the basal membrane of the BBB. We observed a two fold decrease in the expression of Col IV in hPFF injected animals when compared to injection controls suggesting impairment of BBB’s structural integrity (Fig 5A, B). When the BBB is compromised, IgGs are able to cross into the brain parenchyma. Representative images show the extravasation of murine IgG in the SN, Striatum, Cortex and Hippocampus of hPFF-injected mice compared with controls (Fig 5C). Quantitative analysis found a significant increase in murine IgG levels in midbrain of PFF-injected mice compared with saline controls in the ipsilateral hemisphere (Fig 5D). It is unclear when the BBB became disrupted in relation to the PFF injections and synuclein pathology, however by the terminal timepoint of 60 days post-injection, there was a significant positive correlation between pS129 α-Synuclein staining and mIgG extravasation measured in the ipsilateral midbrain (Fig. 5E).

Neuroinflammation in PD is also manifested by activated astrocytes [52]. Moreover, microglial activation and BBB leakage as observed in this study can trigger astrocytic activation wherein astrocytes adhere to a larger cell body and attenuated processes [53]. To test this, we immunostained the ventral midbrain sections with GFAP where phospho Syn pathology was dominant in PFF injected animals. We observed an increase in the astrocyte numbers, indicated by GFAP positive area and increase in GFAP intensity suggesting astrocyte activation in the PFF injected mice midbrain when compared to injection controls (Fig. 5F, G). Taken together, the data here confirms that SN hPFF injection in the A53T mice compromises the BBB integrity, and further suggests one of the plausible reasons behind T cell infiltration in the brain as observed in Fig 4.

## DISCUSSION

Our study demonstrates that injecting human PFFs into the SN of A53T transgenic mice induces bilateral α-Synuclein pathology that spreads from the SN to the striatum, cortex, and other distal regions. Elevated pS129 pathology correlated with markers of neuroinflammation, including microglial and astrocytic activation. T-cell infiltration and BBB compromise were also detected. Significant bilateral dopaminergic neurodegeneration in the SN and striatum was observed and quantified using an AI-based workflow for ROI specific TH intensity analysis. Together, this model provides a promising preclinical methodology for rapidly evaluating therapeutics targeting human α-Synuclein while assessing multiple PD-associated phenotypes within 60 days.

We have previously shown that hPFF injections into the striatum of A53T M83 mice develop time-dependent elevations in pS129 pathology that spreads from the site of injection, striatum, to midbrain and brainstem [25,26] by 60 days post-injection and continues to increase by 90 days, similar to the original findings by Luk et al, 2012 [19]. Striatal injections of hPFFs induced neuroinflammation in brain regions associated with elevated pS129 pathology, but direct measurements of dopaminergic loss was not assessed [25,26]. In this study, we aimed to determine the severity and spread of synuclein pathology following a unilateral injection of hPFFs into the SN, and if this would manifest into greater pathology and associated dopaminergic neuronal degeneration or loss. Indeed, we found that unilateral SN injection of hPFF induced extensive pS129 synuclein pathology in both ipsilateral and contralateral midbrain, dorsal striatum and cortex. pS129 pathology was also found in the olfactory bulb, hippocampus, superior colliculus and amygdala. Importantly, no changes in synuclein pathology was observed in the naive or control injected A53T M83 mice. These findings suggest that the hPFFs seeds can propagate from the site of injection and travel along synaptically connected pathways [54–56], and that the transgene derived α-synuclein does not induce the same level of pS129 pathology without seeding at this age. Studies using direct injections of PFFs into the SN are far fewer than those using striatal injections. In fact, in silico models have been developed predicting the spread of synuclein pathology when injected into the SN [24], but have not been tested experimentally. Our experimental observations match the predictions to some degree, with synuclein spreading from the SN to the dorsal striatum and cortex. However, the degree of spreading to the contralateral brain regions is greater than predicted. Possible explanations are that the model was based on mouse PFFs injections into WT mice, whereas we are using hPFFs injected into A53T M83 transgenic mice. Accelerated premature mortality, body weight loss, and behavioral alterations were observed by 60 days post-nigral injection of hPFFs in A53T M83 mice, whereas striatal injections of hPFFs in M83 mice or mouse PFFs in WT mice take longer to manifest [15,19,25,26,57], suggesting injections into the SN may facilitate a more aggressive phenotype than striatal injections.

Neuroinflammation is a common phenotype observed in many PD mouse models [10,30–32,34,37] Consistent with our prior findings from striatal PFF injections in A53T mice at 90 days post PFF injection [26], we observed an increase in CD68 and GFAP intensity–indicating microglial and astrocytic activation, the ipsilateral midbrain of the PFF injected mice at 60 days post injection. This findings also highlights the aggressive nature of SN PFF injections compared to striatal PFF injection. Although less pronounced, microglial inflammation was also detected on the contralateral side, suggesting that neuroinflammation follows a pattern similar to α-Synuclein pathology progression. The positive correlation between CD68 positive microglial activation and α-Synuclein pathology further supports the role of α-Synuclein aggregates in potentially driving microglial activation in this model. Alternatively, the observed dopaminergic neurodegeneration in the midbrain (Fig 4 B, E) suggests that microglial activation may contribute to the observed neurodegeneration, or be in response to the neurodegeneration [32,34,37].

Studies of post-mortem human PD brains as well as animal models reveal evidence of peripheral immune cell recruitment in response to α-Synuclein aggregation [40,43,45,46,60]. Specifically, T cell infiltration has been studied in both Tg and non-Tg mice following PFF injection, with most studies relying primarily on bulk quantitative assays such as flow cytometry based cell sorting [45–47]. However, one study specifically investigated T cell infiltration in Line 61 Tg mice and observed T cell infiltration in the neocortex and striatum, correlating with neuroinflammation and neurodegeneration. [47]. In our study, we detected T cell infiltration spatially in A53T Tg mice following PFF injection, in the hippocampus, cortex and midbrain, which includes the Substantia Nigra and neighbouring regions. The absence of T cell infiltration in injection-only control Tg mice suggests that this phenotype arises from the interplay of PFF injection and transgene expression, rather than as a result of the injection or transgene alone at this age. Though more prominent on the ipsilateral side, increased T cell infiltration was also observed contralaterally, suggesting that this phenotype is closely tied to the severity of pathology and neuroinflammation at this stage in the PFF-injected mice.

The observed T-cell infiltration in PFF injected mice raises an important question regarding the mechanism by which these immune cells access the CNS parenchyma. One plausible explanation is that T cells traverse into the parenchyma due to BBB leakage. BBB impairment has been reported in PD patients and animal models [61–64] including recent studies in aged human α-Synuclein Tg mice [13,14]. For instance, one study in BAC Tg mice expressing mutant A53T synuclein showed reduced Col IV intensity-an indicator of BBB impairment–in the striatum at 13 months of age [13], while another study reported reduced capillary density in the striatum of Line 61 Tg mice at 6 months of age [14]. However, BBB impairment has not been reported in PFF injection models using WT or SNCA Tg mice. In our study, we observed BBB impairment and leakage in the M83 A53T Tg mice after PFF injection evidenced by reduced ColV expression and IgG extravasation. Notably, IgG extravasation was detected in multiple anatomical regions, aligning with the BBB impairment phenotypes described in prior studies [13] while also extending these findings to the SN. The lack of BBB impairment in injection only controls further indicates that this phenotypic alteration is not a direct consequence of intranigral injections. Additionally, we observed a strong correlation between α-Synuclein pathology and IgG extravasation in the ipsilateral midbrain, suggesting that α-Synuclein pathology may play a key role in driving BBB impairment. Further studies are needed to elucidate the causal relationship between these events.

DA neurodegeneration is a key hallmark of PD, and PFF injection models have reliably shown DA neurodegeneration in both the striatum and the SN [19,23,40]. At 60 days post PFF SN injection, our model demonstrated bilateral neurodegeneration, with approximately 50% reduction in TH intensity in the SN and 38 % in the Striatum. α-Synuclein associated pathology may cause cell autonomous dysfunction leading to this dopaminergic neurodegeneration, or pS129 dependent microglial activation may have triggered inflammatory responses leading to dopaminergic neurodegeneration [31,65]. The bilateral degeneration observed, despite unilateral injection, is likely due to bilateral pS129 SN spread through neuronal SN connectivity, supporting the concept that transhemispheric propagation of pathology induces dopaminergic neurodegeneration [54–56].

PD is a multifaceted neurodegenerative disorder marked by a variety of pathologies, including α-Synuclein pathology, neuroinflammation, T cell infiltration, BBB impairment and dopaminergic neurodegeneration [1,30,35,36,51]. Although numerous animal models of α-Synuclein-related pathology have been developed, none can comprehensively replicate all phenotypes observed in human PD patients. The injection of PFFs directly into the deep brain, as employed in this model, is an artificial method of inducing disease symptoms and must be critically considered when interpreting findings. Additionally, the model relies on a transgenic background with an artificially overexpressed rare synuclein mutation that is not present in the majority of PD patients, further limiting its direct clinical relevance. Despite these limitations, the model’s ability to recapitulate multiple PD-associated phenotypes at a single timepoint underscores its utility. It offers a valuable platform for testing drugs that target human α-Synuclein and assessing their potential to rescue multiple phenotypes within a relatively short timeframe.

## AUTHOR CONTRIBUTIONS

SG: conceptualization, in-vivo study design and execution, AI workflow development, data analysis and writing; SC: in-vivo study execution, data analysis; HL: in-vivo study execution, data analysis HN:Image analysis ; NZ: EM experimental execution and data analysis; RR:Image acquisition ; KS: Tissue management and NSA outsourcing; OF: Histopathology supervision and review; SH: AI workflow development and data-analysis; AE: conceptualization and supervision; BB: conceptualization, supervision and review; WM: conceptualization, supervision and writing.

## ACKNOWLEDGEMENTS

We would also like to thank Neuroscience Associates Inc. (Knoxville, TN, USA) for providing the tissue processing infrastructure to conduct this study.

---

**Supplementary Figure 1.**
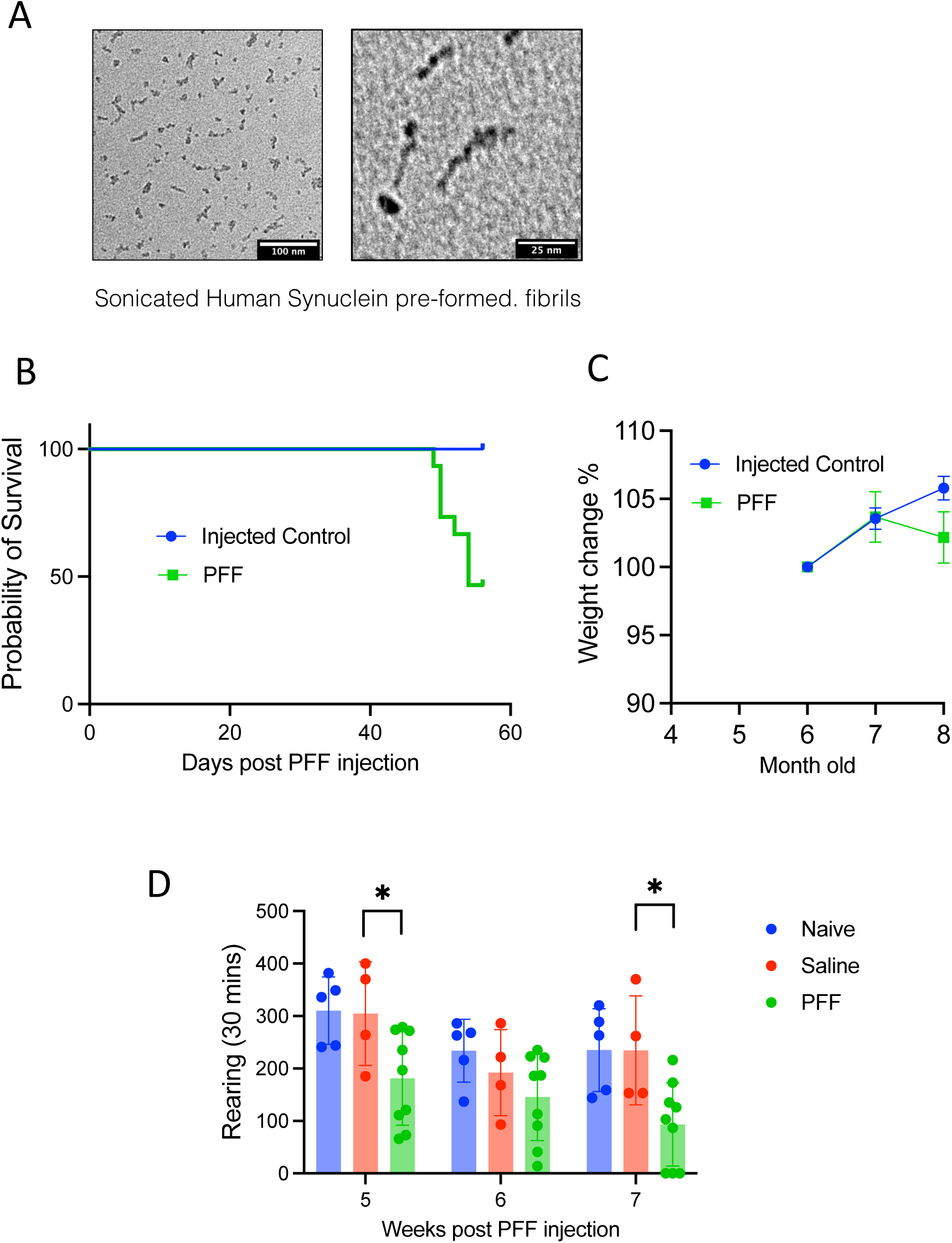
Imaging of PFF fibrils in silico and its effect on survival and weight loss in-vivo.

**Supplementary Figure 2.**
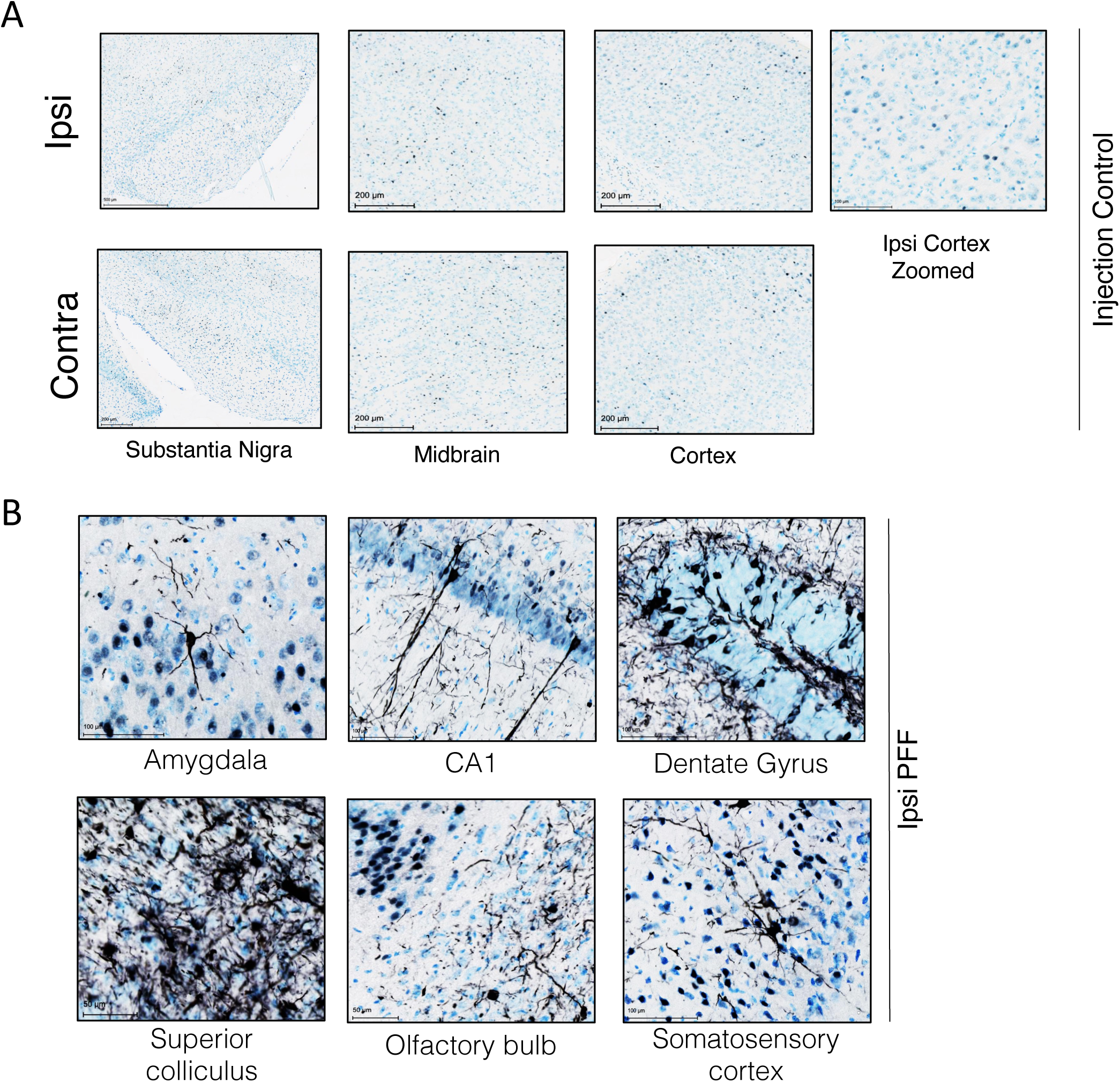
Unilateral PFF injection leads to p-syn associated pathology spreading across different anatomical regions in the brain.

**Supplementary Figure 3.**
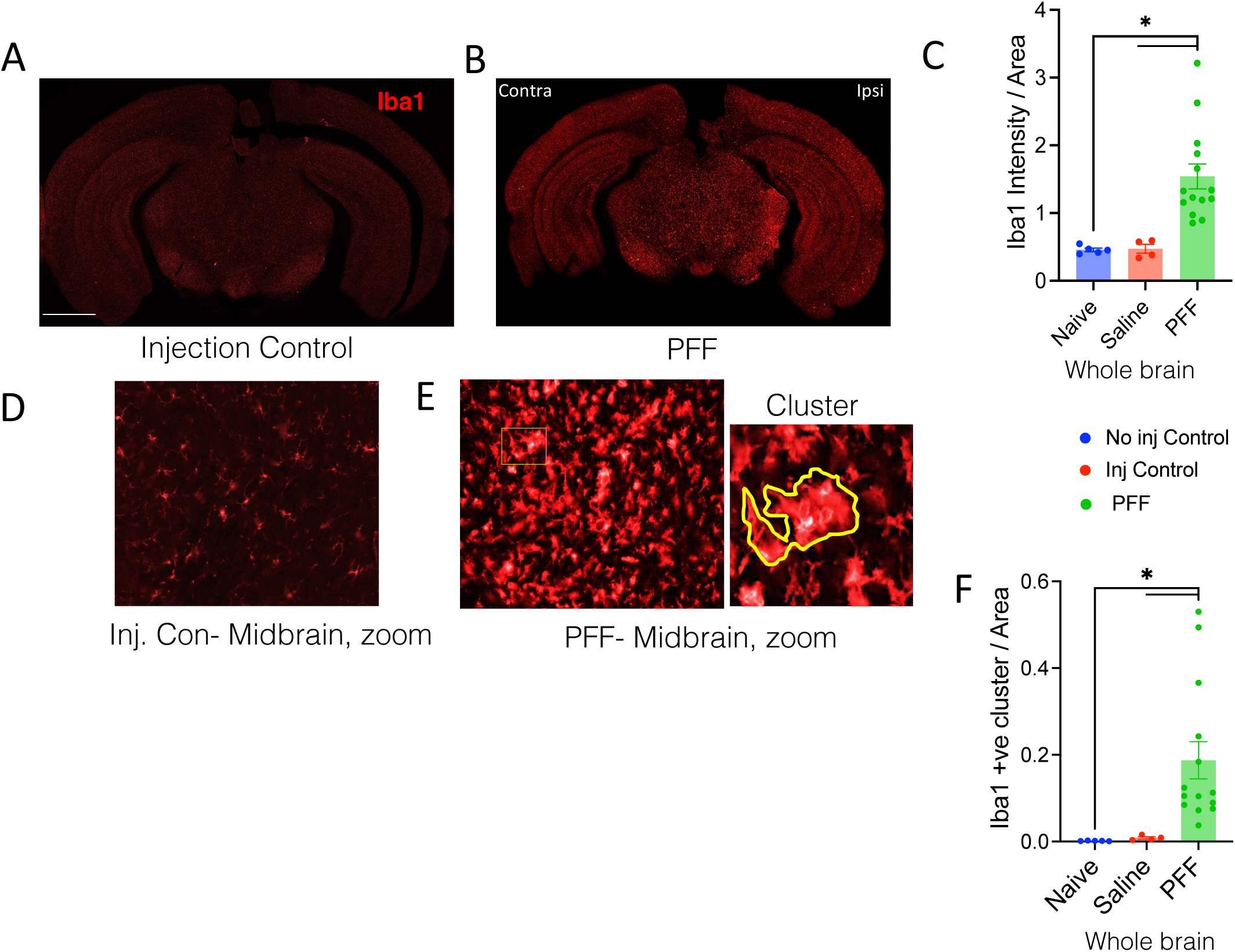
Unilateral PFF injection leads to microglial activation and clustering in midbrain.

**Supplementary Figure 4.**
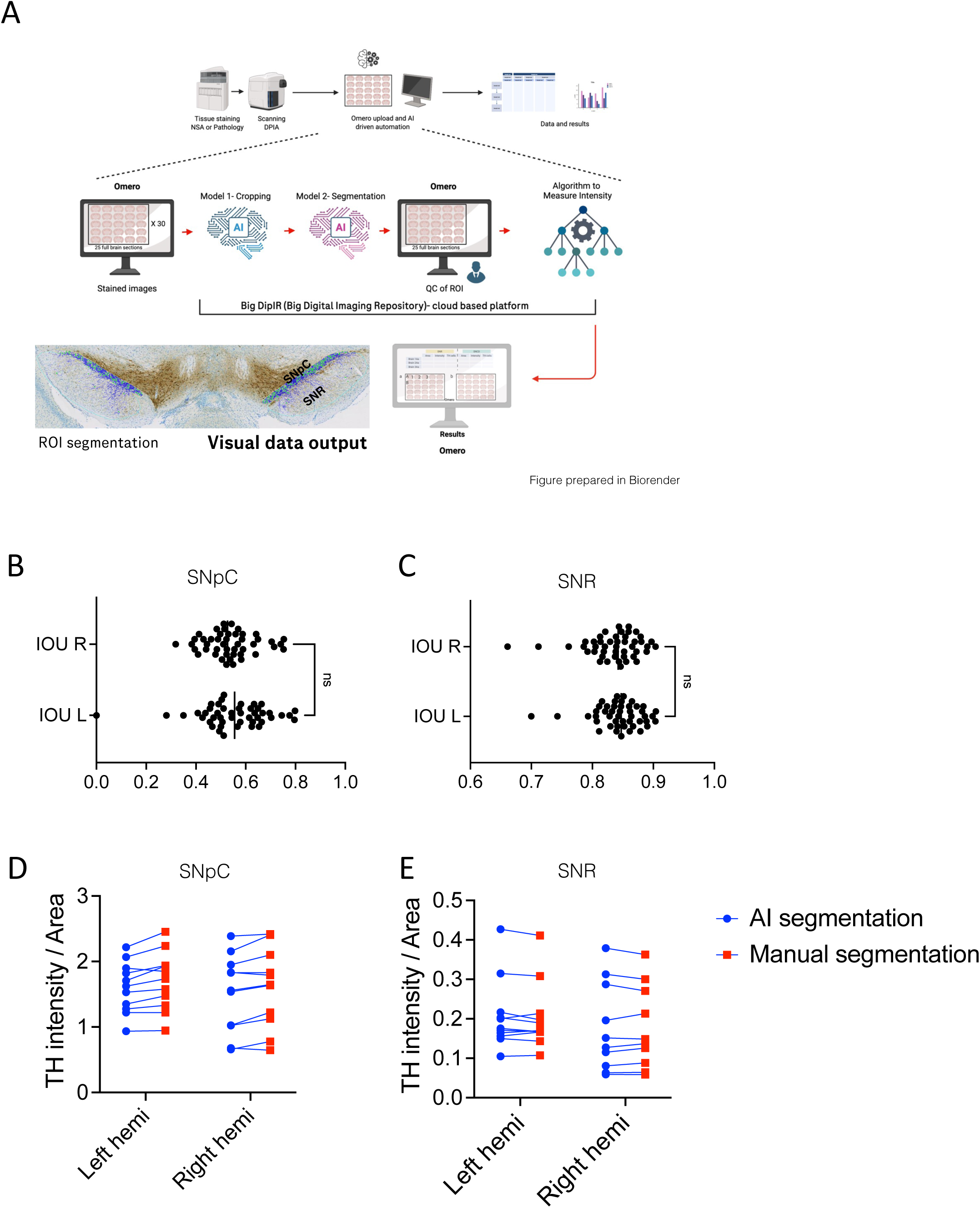
Validation of AI platform used for evaluation of TH status in the substantia Nigra.

**Supplementary Figure 5.**
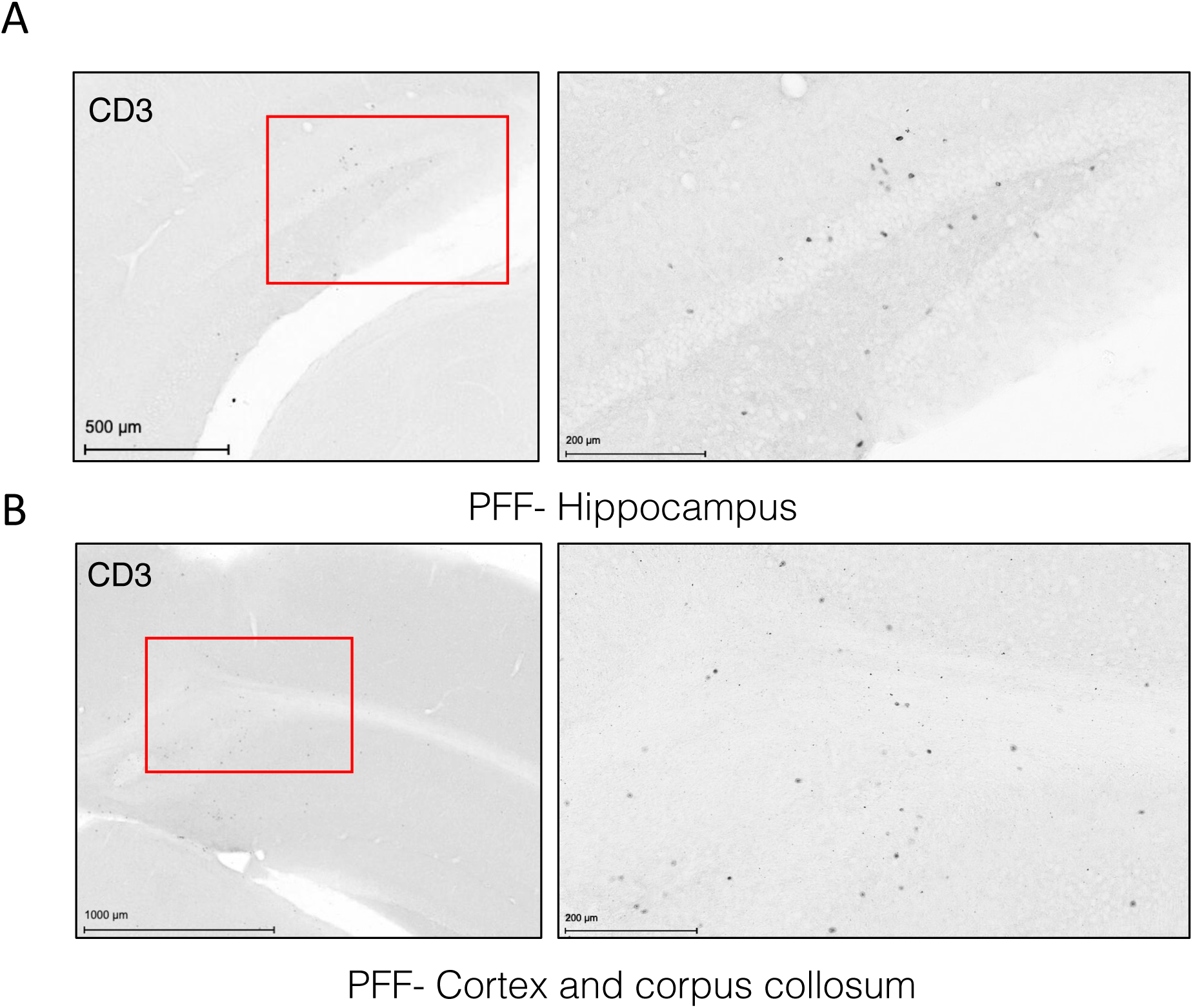
PFF injections lead to T-cell infiltration in the hippocampus and cortex.

