## Supplementary for "Intranigral injection of Alpha-Synuclein pre-formed fibrils leads to BBB compromise and Bilateral Dopaminergic Neurodegeneration in A53T Alpha-Synuclein transgenic mice": Supplementary Material Ghosh 2025 .docx (1).pdf

### Supplementary Figure Legends

**Supplementary Figure 1** Imaging of PFF fibrils in silico and its effect on survival, behavior and weight loss in-vivo **A)** Representative TEM images of sonicated PFF material used for in vivo studies, Scale bar left panel- 100 nm, Scale bar right panel- 25 nm **B)** Survival of saline and PFF injected animals in the study plotted by Kaplan Meier plot. The plot shows survival post injection until the end of the study (8 weeks or 60 days). **C)** Change in body weights (normalized to baseline and weight converted to survival %) between saline and PFF injected animals post PFF or saline injection plotted by survival plot. **C)** Number of rearing spanning over 30 minutes in open field compared between Naïve, Saline injected and PFF injected animals at 5, 6 and 7-weeks post PFF injection. (n= 5 naive, 4 saline, 9 PFF), Kruskal-Wallis test followed by Dunn's multiple comparison test, **p-value <= 0.005. Asterisks (*) denote pairwise statistical significance.

**Supplementary Figure 2** Unilateral PFF injection leads to p-syn associated pathology spreading across different anatomical regions in the brain **A)** Representative images depicting the pS129 positive immunostaining (black) in ipsilateral and contralateral side of saline injected control animals similar to PFF injected animals shown in Fig 1C. SN, Midbrain and Cortex regions are shown here. SN Scale bar: SN- 500 μm, Midbrain and Cortex- 200 μm, Ipsilateral cortex Zoomed- 100 μm. **B)** Representative images depicting the p-syn positive immunostaining (black) in ipsilateral side of PFF injected animals in the follow regions: Amygdala (100 μm), CA1 (100 μm), Dentate Gyrus (100 μm), Superior Colliculus (50 μm), Olfactory Bulb (50 μm) and Somatosensory Cortex (50 μm).

**Supplementary Figure 3** Unilateral PFF injection leads to microglial activation and clustering in midbrain **A & B)** Representative images depicting the IBA1 positive immunofluorescence (red) in saline and PFF injected animals, Scale bar- 2000 μm. Magnified images in the lower panel depict IBA1 positive microglia phenotype and cluster formation in the midbrain. 20X zoom for phenotype and 40 X zoom for cluster **C & D)** Quantification of IBA1 positive signal intensity and IBA1 positive cluster signal intensity normalized to the entire brain section area shown here that covers Midbrain and Cerebral cortex together (n= 5 non-injected, 4 saline, 14 PFF), Kruskal-Wallis test followed by Dunn's multiple comparison test, *p-value > 0.01. Asterisks (*) denote pairwise statistical significance.

**Supplementary Figure 4** Validation of AI platform used for evaluation of TH status in the substantia Nigra **A)** Flowchart showing the workflow for AI powered TH analysis. Specifically, two AI models help crop and measure the TH intensity as the output of the workflow. **B & C)** Quantification of interaction over union (IOU) for the segmented areas by AI in right vs left hemi in the SNR and SNpC of the animals. (n = 46 animals). Max-Whitney t test to test null hypothesis and significance. ns- not significant **D & E)** Graph showing the comparison of output for TH intensity values normalized to area between manual segmentation and AI segmentation for both hemispheres in SNR and SNpC. (n = 11 animals randomly selected from the study).

**Supplementary Figure 5.** PFF injections lead to T-cell infiltration in the hippocampus and cortex. Representative images depicting the CD3 positive T cell immunostaining in the ipsilateral side of the hippocampus (A) and cortex (B) of PFF injected animals. Scale bar- 500 μm-A and 1000 μm-B. Red boxes represent the magnified images for the CD3 positive staining. Scale bar- 200 μm.
